# *APP*-induced patterned neurodegeneration exacerbated by *APOE4* in *C. elegans*

**DOI:** 10.1101/376095

**Authors:** Wisath Sae-Lee, Luisa L. Scott, Aliyah J. Encarnacion, Pragati Kore, Lashaun O. Oyibo, Congxi Ye, Jonathan T. Pierce

**Author notes:** equal contribution (Wisath Sae-Lee, Luisa L. Scott).

## Abstract

Genetic and epidemiological studies have found that variations in the amyloid precursor protein (*APP*) and the apoliopoprotein E (*APOE*) genes represent major modifiers of the progressive neurodegeneration in Alzheimer’s disease (AD). An extra copy or gain-of-function mutations in *APP* lower age of AD onset. Compared to the other isoforms (*APOE3* and *APOE2*), the ε4 allele of *APOE* (*APOE4*) hastens and exacerbates early and late onset forms of AD. Convenient *in vivo* models to study how APP and APOE4 interact at the cellular and molecular level to influence neurodegeneration are lacking. Here, we show that the nematode *C. elegans* can model important aspects of AD including age-related, patterned neurodegeneration that is exacerbated by *APOE4*. Specifically, we found that *APOE4*, but not *APOE3*, acts with *APP* to hasten and expand the pattern of cholinergic neurodegeneration caused by *APP*. Molecular mechanisms underlying how APP and APOE4 synergize to kill some neurons while leaving others unaffected may be uncovered using this convenient worm model of neurodegeneration.

## Introduction

Alzheimer’s disease (AD) is a progressive neurodegenerative disease which cannot be prevented, cured, or decelerated. About 11% of people older than 65 in the US currently suffer from AD (Alzheimer’s Association 2015). Genetic and epidemiological studies have linked variation in several genes strongly to AD. Gain-of-function mutations in amyloid precursor protein (*APP*) or in presenilin genes associated with processing APP are linked to early-onset AD (Goate *et al*. 1991). Possession of extra wild-type copies of *APP*, such as found in Down Syndrome (DS) or in rare individuals without DS, is also linked to early-onset AD (Prasher *et al*. 1998; Cabrejo *et al*. 2006). Additionally, a distinct variant of *APP* appears protective against AD (Jonsson *et al*. 2012). The strongest genetic risk factor for the more common late-onset AD is the ε4 allele of apolipoprotein E (*APOE4*) (Corder *et al*. 1993; Liu *et al*. 2013). Over 65-80% of AD patients carry an *APOE4* allele (Saunders *et al*. 1993; Farrer *et al*. 1997). The life-time risk for AD in people homozygous for *APOE4* is extremely high (91%) relative to those with the more common *APOE3* allele (20%) (Corder *et al*. 1993). *APOE4* is linked to more pronounced neurodegeneration (Holtzman *et al*. 2012); consequently, each copy of *APOE4* decreases the predicted age of death relative to the more common *APOE3* variant (Liu *et al*. 2013). Even for early-onset AD, the *APOE4* allele is associated earlier onset and more severe disease, including elevating levels of the amyloid-β_1-40_ peptide in the case of Down syndrome (Patel *et al*. 2011; Head *et al*. 2011). This suggests that *APP* and *APOE4* may contribute an additive or synergistic risk for AD onset and progression.

Despite the substantial influence of *APOE4* on the progression of AD, how APOE4 modulates the molecular mechanisms underlying neurodegeneration remains elusive. Study of genetically modified mice has contributed to our understanding of the susceptibility to AD and other neurodegenerative diseases conferred by mutations or variants in *APP*, *APOE*, and related genes (Di Battista *et al*. 2016). However, progress towards understanding the mechanistic underpinnings of some AD-related pathologies, particularly neurodegeneration, has been slower (reviewed in LaFerla and Green, 2012). Many mouse models of AD do not show the extensive neuron loss that is present in the human condition. Moreover, it is more expensive and time-consuming to study age-related diseases in mice.

To surmount some of these limitations, we explored whether *APOE* could be studied in the context of *APP*-related neurodegeneration with the genetically tractable model nematode, *Caenorhabditis elegans* (Yi *et al*. 2017). This minimal *in vivo* animal model offers several advantages for basic and applied research for AD. First, worms mature rapidly to adulthood reaching “middle-age” of adulthood only 3 days later as reproduction declines. Despite this compressed life span, *C. elegans* shares many of the genetic, cellular, and molecular processes of aging with humans and mice (Arey and Murphy 2016). Thus, age-dependent processes can be studied within the span of one week with *C. elegans*. Second, forward and reverse genetics as well as transgenesis studies are extremely rapid with *C. elegans*. Third, every cell in the tiny (1 mm) worm is identified, including its 302 neurons, which can be examined individually using fluorescent reporters in the living, transparent worm. The function and health of even single identified neurons can be further probed by quantifying simple behaviors such as egg laying and locomotion. Fourth, several models of neurodegeneration related to AD have been generated using *C. elegans* (Griffin *et al*. 2017). We recently described a transgenic worm strain that expresses a single copy of human *APP* and displays degeneration of a subset of cholinergic neurons in middle-age adulthood (Yi *et al*. 2017; Mondal *et al*. 2018). Using this APP-expressing strain, we discovered small molecules that prevent degeneration of neurons in worm via a conserved signaling pathway and boost cognition in two mouse models of AD (Yi *et al*. 2017). Thus, although the simple worm cannot be used to study the important decline in cognition and memory as in mouse models, it can be used to directly study degeneration of identified neurons and to indirectly predict effective pharmacological approaches in rodent models of AD.

In this study, we tested how expressing variants of human *APOE* altered health and function of neurons in our worm model of APP-related neurodegeneration. We observed several key characteristics associated with AD. First, *APOE4* caused a higher level of neurodegeneration than *APOE3*; and, *APOE4* but not *APOE3* acted in concert with *APP* to further increase neurodegeneration. This retains the isoform-specific effect of *APOE* that is well-documented in human (Corder *et al*. 1993). Second, *APOE4* accelerated neurodegeneration in *C. elegans*, lowering the age of onset from late to early adulthood. This mirrors the expedited onset of degeneration in humans who carry *APOE4* (Holtzman *et al*. 2012). Lastly, despite deliberate expression of *APP* and *APOE4* throughout the nervous system, the pattern of neurodegeneration in the worm model was restricted. This result mimics the fact that earlier and more extensive degeneration is found in specific brain regions, i.e. the hippocampus and the entorhinal cortex of AD patients despite the near-ubiquitous expression of APP and APOE in the human brain (Holtzman *et al*. 2012). Because *APOE* variant expression influences *C. elegans* neurodegeneration in a manner similar to several key aspects found in human AD, our worm model may facilitate both the discovery of molecular pathways involved in the development of AD as well as novel drug discovery for the treatment of AD.

## Materials and Methods

### Plasmid and transgenic animals

The plasmid constructs for human *APP* or *APOE* transgenes under a pan-neuronal promoter (*prab-3::APP::mCherry::UNC-54 UTR; prab-3::APOE4::UNC54 UTR; prab-3::APOE3::UNC54 UTR*) were generated using Multi-site gateway technology (Invitrogen, Carlsbad, CA). Middle entry clones were made using the MultiSite Gateway Three-Fragment Vector Construction Kit (Invitrogen). cDNA for human *APP695* or *APOE* variants (*APOE3, APOE4*) was amplified using the High Fidelity Phusion Polymerase (NEB) and recombined into pDONR221 using the BP clonase (Invitrogen). To generate the expression clones, theses middle entry clones were previously fused into pCFJ150 along with the 5’ *rab-3* promoter and 3’ *unc-54 UTR* entry clones (Yi *et al*. 2014) using the LR clonase II (Invitrogen). Resulting plasmids were verified by sequencing. The construct for the GFP marker for HSN neurons was made by a PCR fusion of a ~2 kb region upstream of *tph-1* amplified from genomic DNA and *GFP::unc-54 UTR* amplified from pPD95.75. The construct for the GFP marker for VA and VB neurons was made by a PCR fusion of a ~1.8 kb region upstream of *del-1* amplified from genomic DNA and *GFP::unc-54 UTR* amplified from pPD95.75. The constructs for expressing *APOE4* in the intestine and coelomocytes were made via NEBuilder^®^ HiFi DNA Assembly. The promoter of *fat-7* (~1.2 kb upstream of the gene) was amplified from genomic DNA while the promoter of *unc-122* (~300 bp) was amplified from pCFJ68.

A single copy of human *APP* under a *rab-3* promoter was inserted into the chromosome II of *C. elegans* via MosSCI transformation (Frøkjær-Jensen *et al*. 2008). The EG4322 strain was selected for the insertion of the construct at ttTi5605 II. Insertion of *APP* was confirmed in this APP strain by PCR and sequencing of the modified region on the chromosome. Extrachromosomal array strains in this study were made by injecting expression plasmids and co-injection markers (1.2 ng/μl for pCFJ90, *pmyo-2:mCherry*, and 30 ng/μl for pCFJ68, *punc-122::GFP, ptph-1::GFP*) described by (Mello *et al*. 1991). The extrachromosomal array for *ptph-1::GFP* was subsequently integrated using standard UV-integration techniques and outcrossed 6 times; this strain served as the wild-type (WT) background for all other strains.

We made three attempts to integrate the extrachromosomal array expressing *APOE4* using traditional UV-integration techniques. However, the attempts were unsuccessful, at least in part due to a low survival rate after UV exposure. Instead, the array [*prab-3::APOE4::UNC54 UTR; pmyo-2:mCherry*] was integrated using CRISPR-cas9-based methods modified from Yoshina, Sawako *et al*. (2016). Briefly, two plasmids containing the *cas9* gene (pDD162) and a guide RNA targeting either the genomic region of integration (LG X:22.84, a location previously used for MosSCI insertions (Frøkjær-Jensen *et al*. 2008)) or a region on the extrachromosomal array (β-lactamase gene), and a co-injection marker (pCFJ104, *pmyo-3:mCherry*) were injected into worms carrying the *APOE4* array. The sequences for guide RNAs used in the integration are as follow, 5’TTAATAGACTGGATGGAGG3’ (β-lactamase gene), 5’ATGTGTCATAAGTCAACAAC3’ and 5’TTATGTAGTCTCTTTCAGTG3’ (X:22.84). The guide RNAs were made according to Ward (2015). After non-homologous repair integrated the linearized array into the chromosome, we chose a strain with a similar intensity of red pharynx as that seen prior to integration.

The integrated *APOE4* line was used to make the APOE4 (JPS844) and APOE4+APP (JPS845) strains. Subsequently, JPS845 was crossed with MT3002 (*ced-3(n1286*) IV) to obtain an APOE4+APP strain with a *ced-3* null background. The genotypes for all the strains used in this study are listed in Table S1.

### Bag of worms assay

Worms were maintained at 20° C as previously described by Brenner (1974). A total of 50 worms were synchronized by picking L4-stage larvae onto NGM agar plates seeded with OP50 bacteria. Worms were carefully examined every day for three days (Day 1-3 of adulthood) for the bag-of-worms phenotype indicated by the presence of hatched larvae within their body. Groups of healthy worms with no hatched larvae were transferred to freshly seeded plate each day. The reported percentage bag-of-worms represents the number of bag-of-worms observed over the period of observation (2 or 3 days) relative to the total number of assayed worms. The assay was repeated 3-4 times for each genotype and expressed as % bag-of-worms ± 95% confidence interval. Group comparisons were made with planned *χ*^2^-tests.

### Scoring of neurodegeneration

Worms maintained at 20° C and synchronized by picking L4-stage larvae onto NGM plates seeded with OP50 bacteria and containing 5-fluoro-2′-deoxyuridine-5′-phosphate (FUDR, 200 μL of a 16.8-mM stock per 12-cm plate) to sterilize adults and prevent non-specific consequences of larvae hatching inside adults. Prior to scoring neuron health, worms were cleaned by transferring them to an unseeded plate until they left no residual tracks of bacteria, a process that took <10 mins. Worms were mounted on 2% agarose pads, immobilized with 30-mM sodium azide and imaged at 40X. For strains with no or extrachromosomal expression of *APOE* (Figure 1), HSN neurons were considered healthy when both HSNL and HSNR were present and the processes were intact. HSNs were considered degenerated when one or both neurons was absent and/or displayed significant morphological abnormalities (e.g, blebbing, beading and absence of processes). Examples of degenerated HSN neurons following these criteria can be found in Figure S1. We observed dimmer GFP expression in the HSNs of integrated *APOE4* strains (Figure 2) making it more difficult to visualize the HSN processes. Thus, HSN health in these strains (JPS844 vs. JPS845) was based exclusively on the presence or absence of the HSN cell-bodies. That is, HSNs were considered to be degenerated when one or both HSN neurons were absent. Neuron health was scored with the observer blind to genotype. An average of 50 worms were scored per strain for each trial. Scoring was repeated 3-4 times for each genotype and expressed as % HSN degeneration ± 95% confidence interval. Group comparisons were made with planned *χ*^2^-tests.

**Figure 1.**
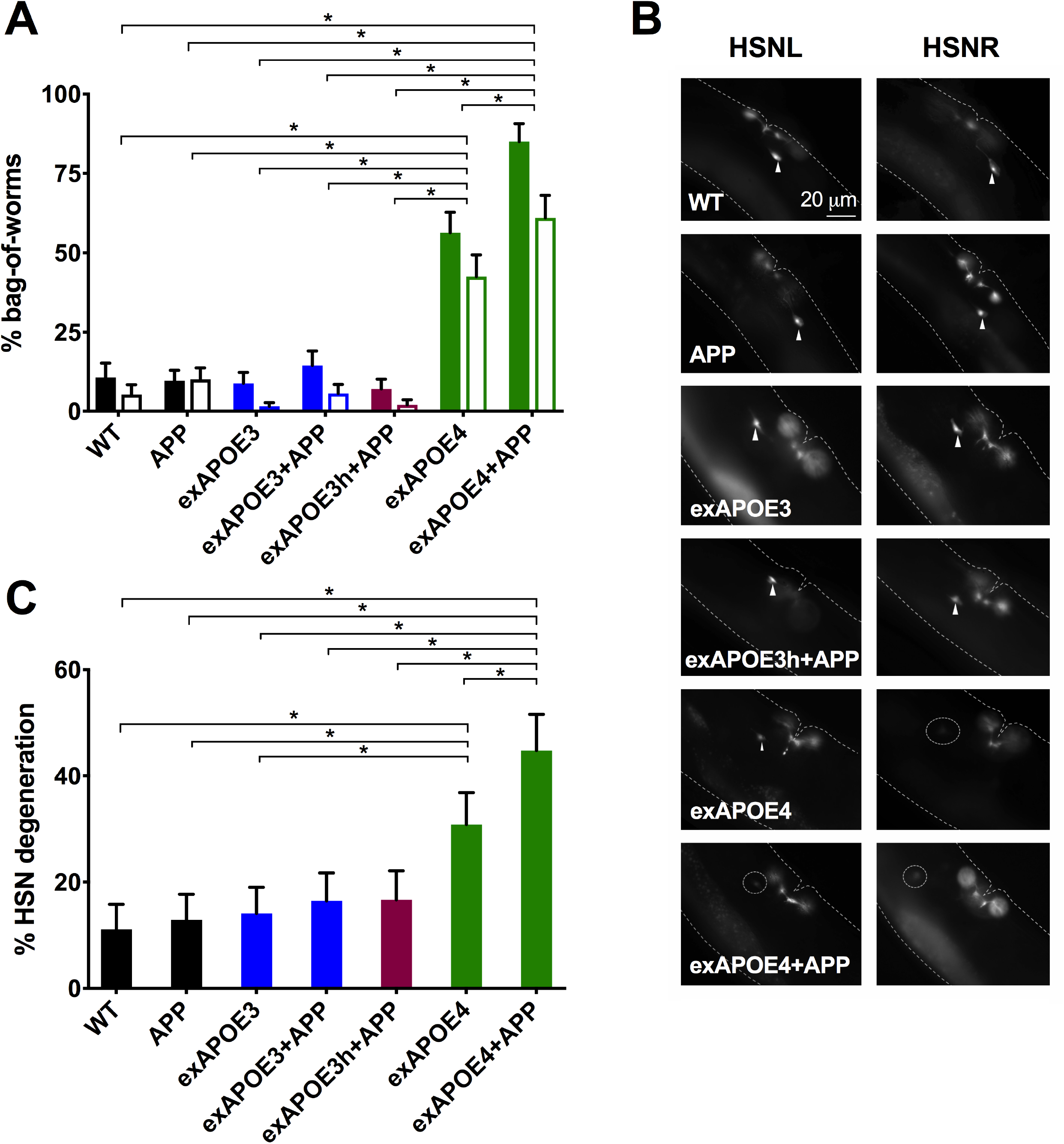
Pan-neuronal expression of *APOE4* induces neurodegeneration in HSN neurons that is exacerbated by concomitant *APP* expression. ***A***, Histogram showing the cumulative percent bag-of-worms phenotype during the first 3 days of adulthood. Bag-of-worms was determined by the presence of progeny with pharyngeal mCherry expression in a hermaphrodite. Two strains of each genotype (signified by grouped pairs of bars) were made with independent extrachromosomal arrays containing a *pmyo-2::mCherry* reporter, and in some cases an *APOE* transgene. The *APP* transgene was integrated. The bag-of-worm phenotype was frequently observed in APOE4-expressing strains with or without *APP* expression but not in strains expressing *APP* alone. In contrast, the expression of *APOE3*, either injected at the same concentration as *APOE4* (exAPOE3) or double the concentration (exAPOE3h), did not increase the frequency of bag-of-worms. For statistical comparisons, shaded bars and open bars were each treated as a set. Within a set, all pairwise comparisons were made with χ^2^ tests. Alpha was set at 0.001 to correct for multiple comparisons (\**p* < 0.001). ***B***, Fluorescent images of the neurons, HSNL and HSNR, in Day 3 adults. Many worms expressing *APOE4* with or without *APP* show morphological abnormalities or a total loss of one or both HSN neurons. Arrowheads indicate healthy HSN neurons. Dotted circles indicate degenerated neurons. C, Histogram showing the percent HSN neurodegeneration on Day 3 of adulthood for a set of strains assayed in A. Pairwise comparisons were made with χ^2^ tests. Alpha was set at 0.003 to correct for multiple comparisons (\**p* < 0.003).

**Figure 2.**
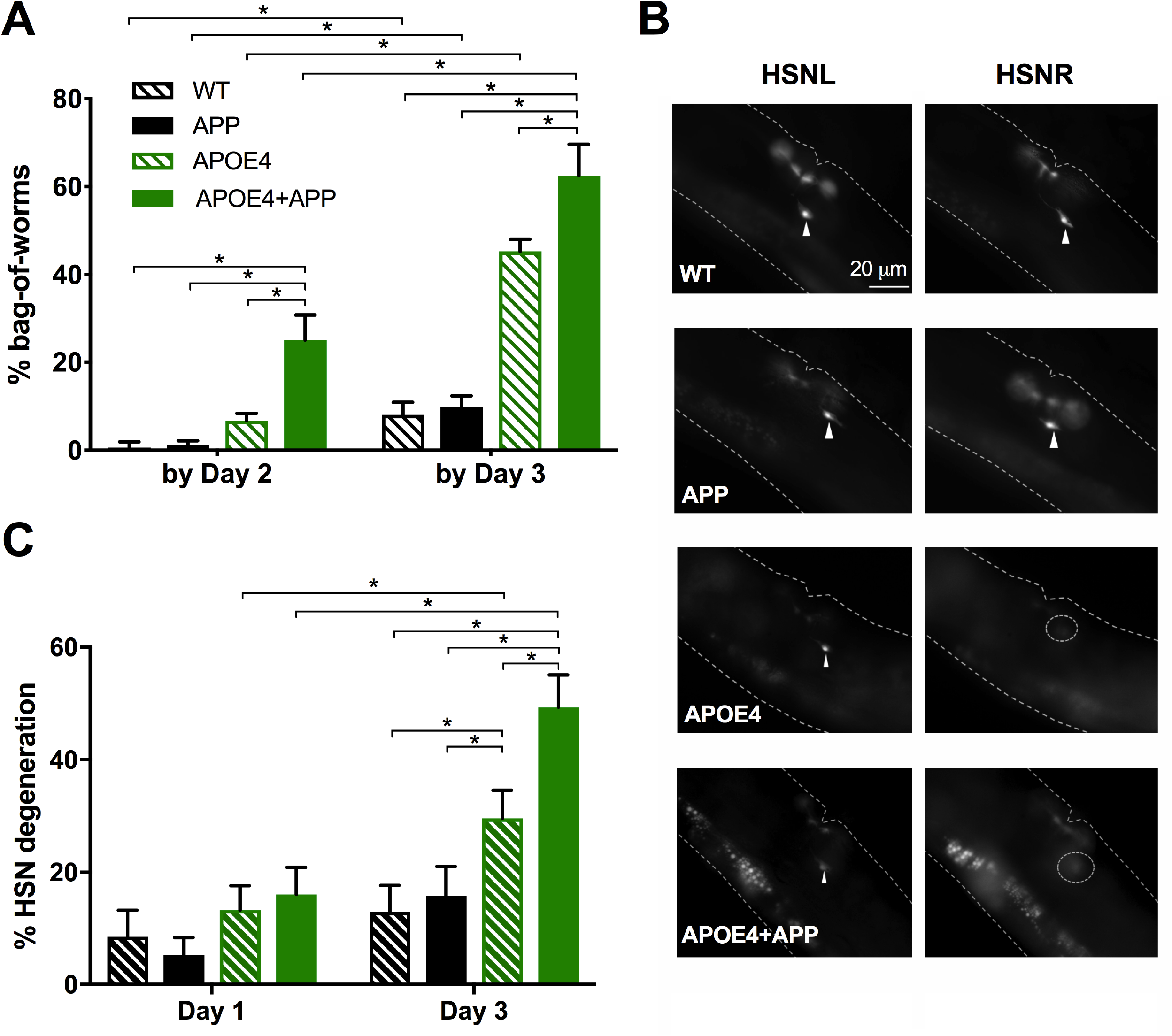
Pan-neuronal co-expression of *APOE4* and *APP* induces age-related neurodegeneration in HSN neurons. ***A***, Histogram showing the cumulative percent bag-of-worms phenotype by Day 2 and Day 3 of adulthood. The frequency of the bag-of-worms phenotype increased at an earlier age for the APOE4+APP strain than for the other genotypes assayed. Both *APP* and *APOE4* transgenes were integrated. ***B***, Fluorescent images of the neurons, HSNL and HSNR, in Day 3 adults. Many APOE4 and APOE4+APP worms show morphological abnormalities or a total loss of one or both HSN neurons. Arrowheads indicate healthy HSN neurons. Dotted circles indicate degenerated neurons. ***C***, Histogram showing the percent HSN neurodegeneration for Day 1 and Day 3 adults. For *A* and *C*, pairwise comparisons within day and within-strain across day were made with χ^2^ tests. Alpha was set at 0.003 to correct for multiple comparisons (\**p* < 0.003).

### Locomotion assay

Worms were cleaned of bacteria as described above. Approximately 15 worms were moved into a 5/8-inch-diameter copper ring sealed on a standard unseeded NGM agar plate. Movement was recorded for 2 min at 2 frames/sec with a FLEA digital camera (Point Gray, Richmond, BC, Canada). The distance that the worms crawled during 1 min was measured using a semiautomated procedure in ImagePro Plus (Media Cybernetics, Rockville, MD) to objectively calculate overall speed of individual worms. Speed for each group is expressed group mean ± SEM. Group comparisons were made with planned Student’s t-tests.

## Results

### *APP* expression exacerbates APOE4-induced neurodegeneration in HSN neurons

We previously developed a *C. elegans* model of neurodegeneration that expressed a single-copy of human *APP* and showed age-related degeneration of a subset of cholinergic neurons (Yi *et al*. 2017; Mondal *et al*. 2018). Specifically, six VC-class neurons located in the middle of the ventral nerve cord begin to die as the worm advances past Day 3 of adulthood. To investigate whether variants in *APOE* modify neurodegeneration phenotypes in our APP worm model, we generated strains that expressed human *APOE4* or *APOE3*, with or without human *APP*. Both *APP* and *APOE* transgenes were expressed throughout the nervous system using the conventional pan-neuronal *rab-3* promoter to mimic widespread brain expression in mammals (Stefanakis *et al*. 2015). *APP* was integrated while initially *APOE* transgenes were expressed with extrachromosomal arrays. Comparisons were made between these strains and a wild-type background (WT).

Despite the degeneration of VC neurons in our APP-expressing strain, the expression of human *APP* alone did not confer any obvious behavioral defects (Mondal *et al*. 2018). To test if *APOE* variants caused gross phenotypic differences that may be indicative of neuronal degeneration or dysfunction beyond that observed in our strain only expressing *APP*, we evaluated the behavior of *APOE3*- or APOE4-expressing strains. We observed an increase in a behavioral phenotype called ‘bag-of-worms’ (BW) in strains expressing *APOE4* but not *APOE3* (Fig. 1A). *C. elegans* normally lays eggs that hatch outside the parent. However, eggs are retained inside the parent under stressful conditions of starvation, or when the neurons that mediate egg-laying become dysfunctional or die (Schafer, 2005). If eggs are retained too long in the worm, they hatch and fill the parent with writhing larvae (Angelo and Van Glist 2009). This BW phenotype can be unambiguously detected at low power via a stereomicroscope and has proven convenient to study many biological processes (e.g. Trent *et al*. 1983; Condradt and Horvitz, 1998).

When we quantified the amount of BW by observing worms for 3 days throughout the *C. elegans* reproductive period, we found that WT worms displayed a low level of BW as expected. The incidence of BW was not raised in strains expressing *APP* or *APOE3*. However, APOE4-expressing strains had a significantly higher percentage of BW than WT, APP-, and APOE3-expressing strains (Fig. 1A). Worms expressing both *APOE4* and *APP* displayed an even higher incidence of BW than worms expressing either *APOE4* or *APP* alone (Fig. 1A). In contrast, the percentage BW in strains expressing both *APOE3* and *APP* remained low and was not significantly higher than that found in strains expressing either gene individually (Fig. 1A). The lack of affect of *APOE3* could not be easily explained by a lower number of *APOE3* transgene copies on the extrachromosome versus *APOE4* copies because an independent strain made with a higher (2x) transformative dose (exAPOE3h) showed a low level of BW indistinguishable from the APP strain (Fig. 1A).

Because the APOE4-expressing strains displayed the BW phenotype even when well fed, this suggested that one or more of the egg-laying neurons were dysfunctional or dying. Egg-laying is controlled primarily by the left-right pair of HSN neurons and to a much lesser extent by VC4 and VC5 neurons (Schafer, 2005). The HSN and VC4 and VC5 neurons connect to each by reciprocal synapses (Altun and Hall, 2018). We checked the health of these neurons by directly visualizing them with a fluorescent reporter through the worm’s transparent body (see methods). We considered HSN neurons “degenerated” when one or both neurons was absent and/or displayed significant morphological abnormalities (e.g, blebbing, beading and absence of processes) (see Fig. 1B and S1). Additionally, we saw clear degeneration of HSN neurons in APOE4-expressing strains with or without co-expression of *APP*. As for BW behavior, the incidence of HSN neuron degeneration was significantly higher in worms expressing both *APP* and *APOE4* than those expressing either *APP* or *APOE4* alone (Fig 1C). Intriguingly, although expression of *APOE4* increased degeneration of HSN neurons, the expression of *APOE3* with or without *APP* co-expression did not increase HSN degeneration relative to WT (Fig. 1C). Even when the *APOE3* transgene was transformed at double the concentration of the *APOE4* transgene and combined with *APP* expression, the resulting strains exhibited a percentage BW and HSN neurodegeneration that was not significantly higher than WT (Fig. 1A,C).

Taken together, these results suggest that the pan-neuronal expression of *APOE4* with *APP* expands degeneration from the VC class neurons to the presynaptic HSN neuron pair. Further, co-expression of *APOE4* with *APP* extends the level of HSN neurodegeneration beyond that observed when either gene is expressed singly.

### Neuronal *APOE4* causes selective neurodegeneration

Thus far, we used extrachromosomal arrays to express the *APOE* transgenes. Although extrachromosomal arrays represent a convenient approach to express transgenes, these arrays are not perfectly carried through cell division which gives rise to mosaic individuals (Leung-Hagesteijn *et al*. 1992). Moreover, the expression of extrachromosomal transgenes are sometimes suppressed (Hsieh and Fire, 2000). To control for these potential caveats, we sought to integrate the *APOE4* transgene array into the worm’s X chromosome and re-evaluate BW phenotype and HSN degeneration. In this way, each cell in the worm would be expected to faithfully express the *APOE4* transgene. We again assayed BW and HSN neurodegeneration, breaking down the incidence as a function of age. The integrated APOE4 and APOE4+APP strains retained a comparable percentage of BW as observed in the extrachromosomal array strains by Day 3 of adulthood (Fig. 1A vs. Fig. 2A). Interestingly, not only was the total incidence of BW higher in the APOE4+APP strain by Day 3 of adulthood, but the total incidence of BW was already substantially raised by Day 2 of adulthood relative to that seen in the APOE4 strain (Fig. 2A). Though both of the APP and APOE4+APP strains showed age-dependent degeneration in HSN neurons, the degree of degeneration was worse in the APOE4+APP strain (Fig. 2C). This is consistent with our findings for the extrachromosomal strains.

Consistent with previous findings, we observed signs of *APP*-related degeneration in the VC4 and VC5 neurons. However, *APOE4* expression greatly exacerbated HSN degeneration but not VC4 and VC5 degeneration when *APOE4* was co-expressed with *APP*. Using our integrated strains, we found that 27% of APOE4+APP worms were missing <u>both</u> HSN neurons on Day 3 of adulthood (vs. 6% in WT, N>300). Less than 1% of either APOE4+APP or WT worms were completely missing VC4 or VC5.

Under the conventional pan-neuronal *rab-3* promoter *APOE4* and *APP* were expressed in almost all neurons; but, aside from the BW phenotype no other abnormal behaviors were grossly apparent in the APOE4+APP strain. This suggests that many neurons in this strain remain intact. For instance, the APOE4+APP strain did not seem to have defects in locomotion compared to WT worms as measured by their speed (Fig. 3B). The crawling speed was indistinguishable on both Day 1 and Day 3 of adulthood between WT and APOE4+APP worms (Fig. 3B). Consistent with this behavioral observation, when we looked at VA and VB cholinergic motor neurons, which control locomotion, and we saw no qualitatively discernable differences in morphology and number of VA and VB neurons between WT and APOE4+APP worms (Fig. 3A).

**Figure 3.**
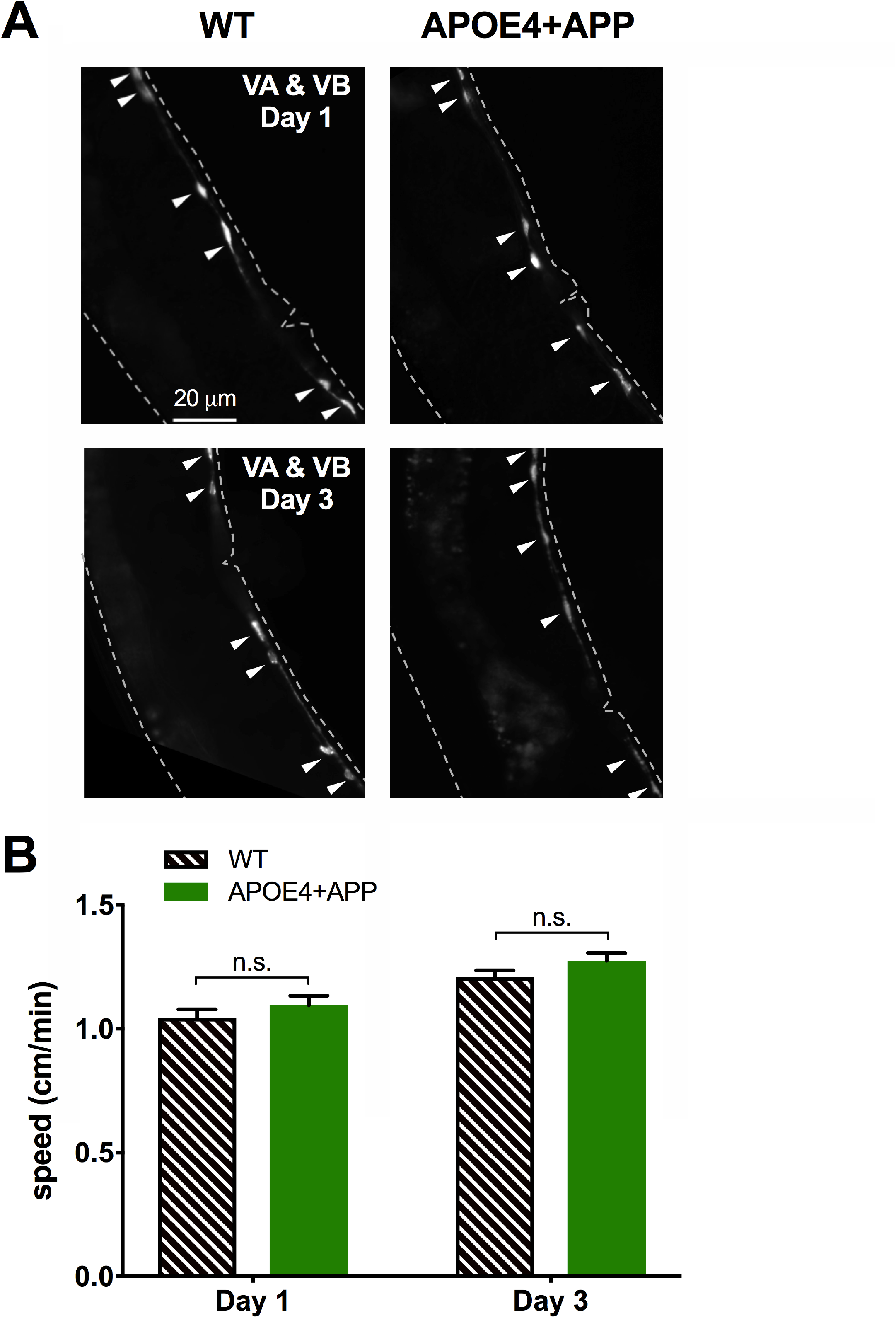
Neurodegeneration induced by pan-neuronal co-expression of *APOE4* and *APP* spares VA and VB neurons. ***A***, Fluorescent images of the VA and VB neurons show no morphological abnormalities in Day 3 APOE4+APP adults relative to Day 1 APOE4+APP or WT adults. Arrowheads indicate healthy VA and VB neurons. Both *APP* and *APOE4* transgenes were integrated. ***B***, Locomotion, a behavior mediated in part by VA and VB neurons, is not disrupted in APOE4+APP worms compared to WT worms. Histogram showing mean crawl speed ± SEM for WT and APOE4+APP worms on Day 1 and Day 3 of adulthood. Strain comparisons were made using Student’s *t*-tests.

Next, we asked whether *APOE4* induces neurodegeneration when expressed outside the nervous system on a pan-neuronal *APP* background. Using extrachromosomal arrays, we expressed *APOE4* in organs involved in metabolic and excretory functions, the coelomocytes and intestine, using the *unc-122* and *fat-7* promoter, respectively. Unlike with pan-neuronally expressed *APOE4*, the incidence of BW was as low for strains expressing *APOE4* in either coelomocytes or intestine as in the background *APP* strains (Fig. 4A). This low percentage of BW for strains with *APOE4* expressed in coelomocytes or intestine corresponded with low levels of HSN degeneration in these strains (Fig. 4B,C).

**Figure 4.**
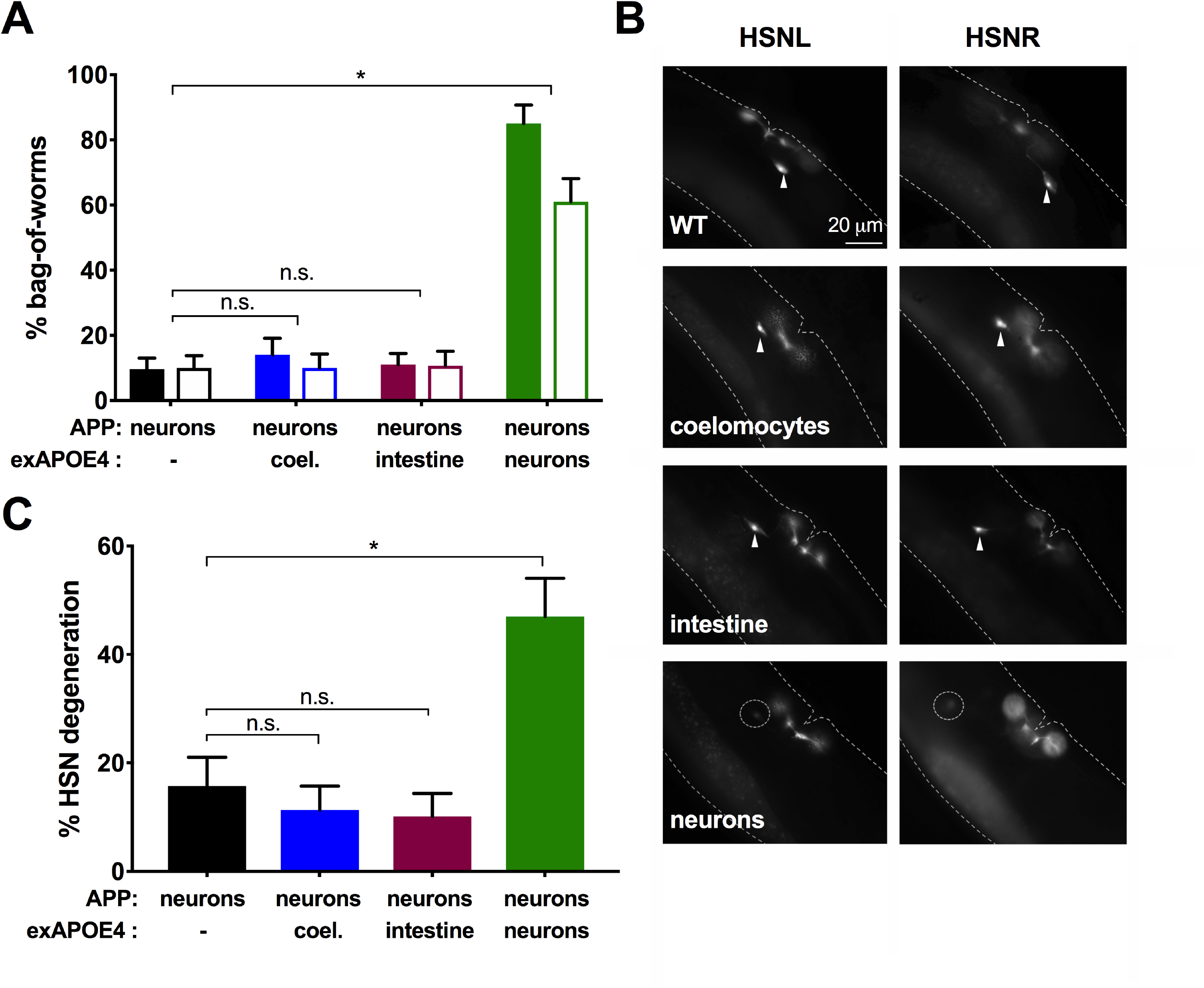
Pan-neuronal, but not non-neuronal, expression of *APOE4* promotes HSN neurodegeneration on a pan-neuronal *APP* background. ***A***, Histogram showing the cumulative percent bag-of-worms phenotype by Day 3 of adulthood. The frequency of the bag-of-worms phenotype increased in worms with pan-neuronal *APP* expression when *APOE4* was expressed in neurons, but not in coelomocytes or intestine. Two strains of each genotype were made (signified by groups of paired bars) with independent extrachromosomal arrays containing a *pmyo-2::mCherry* and *APOE4* transgene. The *APP* transgene was integrated. For statistical comparisons, shaded bars and open bars were each treated as a set. Within a set, each exAPOE4 strain was compared with the background APP strain using χ^2^ tests. Alpha was set at 0.008 to correct for multiple comparisons (\**p* < 0.008). ***B***, Fluorescent images of the neurons, HSNL and HSNR, in Day 3 adults. Many worms expressing pan-neuronal *APOE4* on the *APP* background show morphological abnormalities or a total loss of one or both HSN neurons. Label indicates the location of *APOE4* expression. Arrowheads indicate healthy HSN neurons. Dotted circles indicate degenerated neurons. ***C***, Histogram showing the percent HSN neurodegeneration on Day 3 of adulthood for a set of strains assayed in A. Each exAPOE4 strain was compared with the background APP strain using χ^2^ tests. Alpha was set at 0.015 to correct for multiple comparisons (\**p* < 0.015).

To better understand the mechanism of degeneration, we investigated whether HSN neurons degenerate in manner dependent on apoptosis. We crossed the *ced-3*(*n1286*) null mutation into the integrated APOE4+APP strain. CED-3 is an executioner caspase that is part of the core apoptotic machinery in worm (Yuan *et al*. 1993). We found that the incidence of both BW and HSN neurodegeneration did not decrease in APOE4+APP strains with or without a mutation in this key apoptotic caspase (Fig. 5A-C). Thus, pan-neuronal co-expression of *APOE4* and *APP* appears to cause HSN neurons to degenerate via a mechanism other than apoptosis.

**Figure 5.**
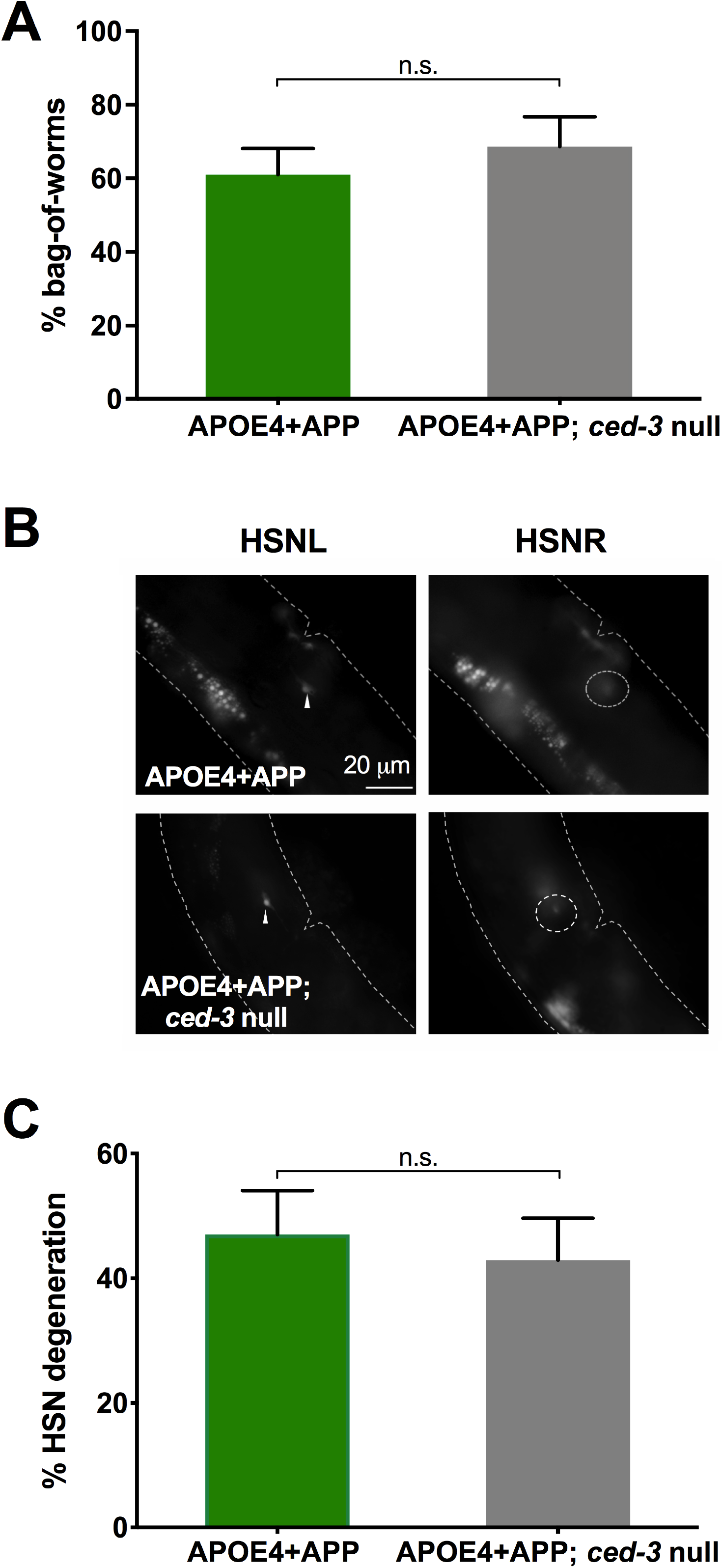
Neurodegeneration induced by pan-neuronal co-expression of *APOE4* and *APP* is not mediated by CED-3. ***A***, Histogram showing the cumulative percent bag-of-worms phenotype by day 3 of adulthood. The frequency of the bag-of-worms phenotype in the APOE4+APP strain is not altered in a *ced-3* null background. Both *APP* and *APOE4* transgenes were integrated. ***B***, Likewise, *ced-3* expression does not alter neurodegeneration in the APOE4+APP strain. Fluorescent images of HSNL and HSNR. Arrowheads indicate healthy HSN neurons. Dotted circles indicate degenerated neurons. ***C***, Histogram showing the percent HSN neurodegeneration for Day 3 adults. Comparisons were made with χ^2^ tests.

## Discussion

In this study, we present a novel Alzheimer’s disease model in *C. elegans*. Although worms cannot model all aspects of human AD, our model mirrors three important characteristics: isoform-specific degeneration, age-dependent degeneration, and cell-specific degeneration. In humans, the allele ε4 of *APOE* (*APOE4*) increases the risk as well as the mean age of clinical onset of AD in a copy-dependent manner (Corder *et al*. 1993). The more common *APOE* variant in humans, *APOE3*, does not confer this relative risk. Our data show that *APOE4*, but not *APOE3*, causes degeneration in a subset of neurons in *C. elegans. APP* co-expression enhances this APOE4-related degeneration of the HSN neurons in *C. elegans*. By contrast, the HSN neurons do not appear to degenerate when *APP* is expressed alone. Similarly, in APOE-target replacement (TR) mouse models engineered on an amyloid-β transgenic background, pathological phenotypes, including amyloid-β deposition, are observed at a higher level in APOE4-TR mice than APOE3-TR mice (Bales *et al*. 2009; Fryer *et al*. 2005). However, studies that show a direct mechanistic link between *APP* and *APOE4* for neurodegeneration are still lacking.

In our worm *APOE4-expressing* strains, HSN neurons display neurodegeneration by becoming dim, shrinking, forming blebs, and disappearing in many animals. We also found that the long axonal processes of HSN neurons, which extend anteriorly to synapse onto neurons in the nerve ring, undergo beading. This is reminiscent of how neuronal processes degenerate after laser axonomy (Yanik *et al*. 2006). In mice, targeted replacement of ApoE with human APOE4 causes dendrites of neurons in the amygdala to shrink compared to the dendrites of APOE3 TR mice (Klein *et al*. 2010). These morphological changes in the processes of dying neurons suggest that there might be conserved pathways for degeneration at this cellular level.

One of the prominent characteristics of AD is age-dependent neurodegeneration. Coexpression of *APP* with *APOE4* throughout the nervous system in *C. elegans* enhances the incidence of APOE4-related neurodegeneration. Interestingly, the cumulative effect of *APP* and *APOE4* expression manifests more strongly in “middle age” adulthood, suggesting that the additive insult may arise from an age-related process. In patients with Down syndrome who carry an extra copy of *APP*, *APOE4* appears to expedite the onset of their early-onset AD and accelerate the progression of neurodegeneration (Patel *et al*. 2011; Head *et al*. 2011).

In humans, neurodegeneration occurs more extensively in certain regions of the brain, such as hippocampus and entorhinal cortex (Saxena and Caroni 2011; Colin *et al*. 2015). This aspect is one of the more curious observations in AD patients because APP and APOE4 are both expressed throughout the brain (Huang and Mucke 2012). Similarly, our results show that the VA and VB cholinergic neurons did not degenerate despite the pan-neuronal expression of *APP* and *APOE4* transgenes and proximity to the vulnerable VC neurons. The restricted pattern of *APOE4*-dependent degeneration in worm suggests that there are cellular and/or molecular characteristics that confer resistance or vulnerability to *APOE4* expression that may make them resistant (e.g., VA and VB) or vulnerable (e.g., HSNs).

Although APOE4 is expressed in both the liver and the nervous system in human (Holtzman *et al*. 2012; Liu *et al*. 2013; Mahley *et al*. 2006), it is unclear whether hepatic, neuronal, or both types of APOE4 contribute to neurodegeneration. We were interested in exploring whether *APOE4*-induced degeneration is tissue-specific. Recent studies have demonstrated clear examples of cell non-autonomous signaling between neuronal and nonneuronal tissue in *C. elegans* (van Oosten-Hawle *et al*. 2013; Taylor and Dillin 2013; Melentijevic *et al*. 2017). For instance, unfolded protein responses, which may underlie neurodegeneration, are interdependently co-activated between the nervous systems and the intestine (van Oosten-Hawle *et al*. 2013; Taylor and Dillin 2013). Additionally, when under stress worm neurons jettison particles extracellularly to become absorbed by coelomocytes (Melentijevic *et al*. 2017). Taken together, these studies suggest that *APOE4* could very well influence neurodegeneration when expressed outside of the neurons. However, after probing expression in both intestine and coelmocytes, our results indicate that *APOE4* mainly induces degeneration when expressed in the nervous system of *C. elegans*. These results are consistent with other work showing the sufficiency of neuronal *APOE4* expression in causing exacerbated cellular or behavioral deficits. For example, in mice neuronal *APP* can cause degeneration and memory deficits when co-expressed with *APOE4* (Bien-Ly *et al*. 2011; Harris *et al*. 2013).

To begin to understand the basis for neurodegeneration in our model, we asked whether degeneration of HSN neurons required apoptosis. CED-3 is the primary executioner caspase that initiates the cascade of apoptosis (Yuan *et al*. 1993). When a *ced-3* null mutation was crossed into the APOE4+APP strain, we saw no reduction of the bag-of-worms phenotype or HSN neurodegeneration. This indicates that apoptosis is likely not involved in *APOE4-* dependent degeneration in *C. elegans*. Other *C. elegans* neurodegenerative models similarly show that *ced-3* and other broad apoptotic mechanisms do not play a role in degeneration (Liachko *et al*. 2010). Despite extensive research into its developmental role, the role of apoptosis in adult *C. elegans* is inconclusive at best. Further studies are required to elucidate the mechanism of neurodegeneration in our APOE4 model in *C. elegans*. Our model is well positioned to be used to explore how patterned neurodegeneration may arise in AD and other neurodegenerative diseases.

---

**Figure S1**. Representative images of degenerating HSN neurons. **A-C**, Degenerated HSN neurons had dim (arrow heads) or barely visible (dotted circle) cell bodies. **D-G**, Earlier evidence of HSN neuron degeneration was visible as blebbing of the cell body (arrows). **H-I**, Processes of degenerating HSN neurons also showed beading (H, arrows) that was not visible in healthy HSN neurons (I). Autofluorescence from intestine bright in panels H and I due to higher gain to visualize HSN axon.

## References

Altun, Z.F. and Hall, D.H. 2018. Reproductive system. In WormAtlas. doi:10.3908/wormatlas.1.17

Alzheimer’s Association. 2015 Alzheimer’s Disease Facts and Figures. Alzheimer’s & Dementia 2015;11(3)332.

Angelo G, Van Gilst MR. 2009. Starvation protects germline stem cells and extends reproductive longevity in C. elegans. Science. 2009 Nov 13;326(5955):954–8. doi: 10.1126/science.1178343. PMID: 19713489

Arey RN, Murphy CT. 2017. Conserved regulators of cognitive aging: From worms to humans. Behav Brain Res. 2017 Mar 30;322(Pt B):299–310. doi: 10.1016/j.bbr.2016.06.035. Epub 2016 Jun 18. PMID: 27329151

Bales KR, Liu F, Wu S, Lin S, Koger D, DeLong C, Hansen JC, Sullivan PM, Paul SM. 2009. Human APOE isoform-dependent effects on brain beta-amyloid levels in PDAPP transgenic mice. J Neurosci. 2009 May 27;29(21):6771–9. doi: 10.1523/JNEUROSCI.0887-09.2009. PMID: 19474305

Bien-Ly N, Andrews-Zwilling Y, Xu Q, Bernardo A, Wang C, Huang Y. 2011. C-terminal-truncated apolipoprotein (apo) E4 inefficiently clears amyloid-beta (Abeta) and acts in concert with Abeta to elicit neuronal and behavioral deficits in mice. Proc Natl Acad Sci U S A. 2011 Mar 8;108(10):4236–41. doi: 10.1073/pnas.1018381108. Epub 2011 Feb 22. PMID: 21368138

Cabrejo L, Guyant-Maréchal L, Laquerrière A, Vercelletto M, De la Fournière F, Thomas-Antérion C, Verny C, Letournel F, Pasquier F, Vital A, Checler F, Frebourg T, Campion D, Hannequin D. 2006. Phenotype associated with APP duplication in five families. Brain. 2006 Nov;129(Pt 11):2966–76. Epub 2006 Sep 7. PMID: 16959815

Conradt B, Horvitz HR. 1998. The C. elegans protein EGL-1 is required for programmed cell death and interacts with the Bcl-2-like protein CED-9. Cell. 1998 May 15;93(4):519–29. PMID: 9604928

Corder EH, Saunders AM, Strittmatter WJ, Schmechel DE, Gaskell PC, Small GW, Roses AD, Haines JL, Pericak-Vance MA.1993. Gene dose of apolipoprotein E type 4 allele and the risk of Alzheimer’s disease in late onset families. Science. 1993 Aug 13;261(5123):921–3. PMID: 8346443

Di Battista AM, Heinsinger NM, Rebeck GW. 2016. Alzheimer’s Disease Genetic Risk Factor APOE-ε4 Also Affects Normal Brain Function. Curr Alzheimer Res. 2016;13(11):1200–1207. Review. PMID: 27033053

Farrer LA, Cupples LA, Haines JL, Hyman B, Kukull WA, Mayeux R, Myers RH, Pericak-Vance MA, Risch N, van Duijn CM. 1997 Effects of age, sex, and ethnicity on the association between apolipoprotein E genotype and Alzheimer disease. A meta-analysis. APOE and Alzheimer Disease Meta Analysis Consortium. JAMA. 1997 Oct 22-29;278(16):1349–56. PMID: 9343467

Frøkjaer-Jensen C, Davis MW, Hopkins CE, Newman BJ, Thummel JM, Olesen SP, Grunnet M, Jorgensen EM. 2008. Single-copy insertion of transgenes in Caenorhabditis elegans. Nat Genet. 2008 Nov;40(11):1375–83. doi: 10.1038/ng.248. Epub 2008 Oct 26. PMID: 18953339

Fryer JD, Simmons K, Parsadanian M, Bales KR, Paul SM, Sullivan PM, Holtzman DM. 2005. Human apolipoprotein E4 alters the amyloid-beta 40:42 ratio and promotes the formation of cerebral amyloid angiopathy in an amyloid precursor protein transgenic model. J Neurosci. 2005 Mar 16;25(11):2803–10. PMID: 15772340

Goate A, Chartier-Harlin MC, Mullan M, Brown J, Crawford F, Fidani L, Giuffra L, Haynes A, Irving N, James L, et al. 1991. Segregation of a missense mutation in the amyloid precursor protein gene with familial Alzheimer’s disease. Nature. 1991 Feb 21;349(6311):704–6. PMID: 1671712

Griffin EF, Caldwell KA, Caldwell GA. Genetic and Pharmacological Discovery for Alzheimer’s Disease Using *Caenorhabditis elegans*. ACS Chem Neurosci. 2017 Dec 20;8(12):2596–2606. doi: 10.1021/acschemneuro.7b00361. Epub 2017 Oct 25. PMID: 29022701

Harris FM, Brecht WJ, Xu Q, Tesseur I, Kekonius L, Wyss-Coray T, Fish JD, Masliah E, Hopkins PC, Scearce-Levie K, Weisgraber KH, Mucke L, Mahley RW, Huang Y. 2003. Carboxyl-terminal-truncated apolipoprotein E4 causes Alzheimer’s disease-like neurodegeneration and behavioral deficits in transgenic mice. Proc Natl Acad Sci U S A. 2003 Sep 16;100(19):10966–71. Epub 2003 Aug 25.PMID: 12939405

Holtzman D. M. and Fagan A. M. (1998) Potential role of apoE in structural plasticity in the nervous system. Implications for disorders of the central nervous system. Trends Cardiovasc. Med. 8, 250–255.

Holtzman, David M., Joachim Herz, and Guojun Bu. “Apolipoprotein E and apolipoprotein E receptors: normal biology and roles in Alzheimer disease.” Cold Spring Harbor perspectives in medicine 2.3 (2012): a006312.

Hsieh J, Fire A. 2000. Recognition and silencing of repeated DNA. Annu Rev Genet. 2000;34:187–204. PMID: 11092826

Huang Y, Mucke L. 2012. Alzheimer mechanisms and therapeutic strategies. Cell. 2012 Mar 16;148(6):1204–22. doi: 10.1016/j.cell.2012.02.040. PMID: 22424230

Jonsson T, Atwal JK, Steinberg S, Snaedal J, Jonsson PV, Bjornsson S, Stefansson H, Sulem P, Gudbjartsson D, Maloney J, Hoyte K, Gustafson A, Liu Y, Lu Y, Bhangale T, Graham RR, Huttenlocher J, Bjornsdottir G, Andreassen OA, Jönsson EG, Palotie A, Behrens TW, Magnusson OT, Kong A, Thorsteinsdottir U, Watts RJ, Stefansson K. 2012. A mutation in APP protects against Alzheimer’s disease and age-related cognitive decline. Nature. 2012 Aug 2;488(7409):96–9. doi: 10.1038/nature11283. PMID: 22801501

Klein RC, Mace BE, Moore SD, Sullivan PM. 2010. Progressive loss of synaptic integrity in human apolipoprotein E4 targeted replacement mice and attenuation by apolipoprotein E2. Neuroscience. 2010 Dec 29;171(4):1265–72. doi: 10.1016/j.neuroscience.2010.10.027. Epub 2010 Oct 15. PMID: 20951774

LaFerla, F. M., and Green, K. N. (2012). Animal Models of Alzheimer Disease. Cold Spring Harbor Perspectives in Medicine, 2(11), a006320. http://doi.org/10.1101/cshperspect.a006320

Leung-Hagesteijn C, Spence AM, Stern BD, Zhou Y, Su MW, Hedgecock EM, Culotti JG. 1992. UNC-5, a transmembrane protein with immunoglobulin and thrombospondin type 1 domains, guides cell and pioneer axon migrations in C. elegans. Cell. 1992 Oct 16;71(2):289–99. PMID: 1384987

Liachko NF, Guthrie CR, Kraemer BC. 2010. Phosphorylation promotes neurotoxicity in a Caenorhabditis elegans model of TDP-43 proteinopathy. J Neurosci. 2010 Dec 1;30(48):16208–19. doi: 10.1523/JNEUROSCI.2911-10.2010. PMID: 21123567

Liu CC, Liu CC, Kanekiyo T, Xu H, Bu G. 2013. Apolipoprotein E and Alzheimer disease: risk, mechanisms and therapy. Nat Rev Neurol. 2013 Feb;9(2):106–18. PMID: 23296339

Mahley RW, Weisgraber KH, Huang Y. 2006. Apolipoprotein E4: a causative factor and therapeutic target in neuropathology, including Alzheimer’s disease. Proc Natl Acad Sci U S A. 2006 Apr 11;103(15):5644–51. Epub 2006 Mar 27. PMID: 16567625

Masters CL, Bateman R, Blennow K, Rowe CC, Sperling RA, Cummings JL. 2015. Alzheimer’s disease. Nat Rev Dis Primers. 2015 Oct 15;1:15056. doi: 10.1038/nrdp.2015.56. PMID: 27188934

Melentijevic I, Toth ML, Arnold ML, Guasp RJ, Harinath G, Nguyen KC, Taub D, Parker JA, Neri C, Gabel CV, Hall DH, Driscoll M. 2017. C. elegans neurons jettison protein aggregates and mitochondria under neurotoxic stress. Nature. 2017 Feb 16;542(7641):367–371. doi: 10.1038/nature21362. Epub 2017 Feb 8. PMID: 28178240

Mello C. C., Kramer J. M., Stinchcomb D., Ambros V., 1991. Efficient gene transfer in C. elegans: extrachromosomal maintenance and integration of transforming sequences. EMBO J. 10: 3959–3970

Mondal S, Hegarty E, Sahn JJ, Scott LL, Gökçe SK, Martin C, Ghorashian N, Satarasinghe PN, Iyer S, Sae-Lee W, Hodges TR, Pierce JT, Martin SF, Ben-Yakar A. High-Content Microfluidic Screening Platform Used To Identify σ2R/Tmem97 Binding Ligands that Reduce Age-Dependent Neurodegeneration in *C. elegans* SC_APP Model. ACS Chem Neurosci. 2018 Feb 23. doi: 10.1021/acschemneuro.7b00428. PMID: 29426225

Patel A, Rees SD, Kelly MA, Bain SC, Barnett AH, Thalitaya D, Prasher VP. 2011. Association of variants within APOE, SORL1, RUNX1, BACE1 and ALDH18A1 with dementia in Alzheimer’s disease in subjects with Down syndrome. Neurosci Lett. 2011 Jan 7;487(2):144–8. doi: 10.1016/j.neulet.2010.10.010. Epub 2010 Oct 12. PMID: 20946940

Saunders AM, Strittmatter WJ, Schmechel D, George-Hyslop PH, Pericak-Vance MA, Joo SH, Rosi BL, Gusella JF, Crapper-MacLachlan DR, Alberts MJ, et al. 1993 Association of apolipoprotein E allele epsilon 4 with late-onset familial and sporadic Alzheimer’s disease. Neurology. 1993 Aug;43(8):1467–72. PMID: 8350998

Saxena S, Caroni P. 2011. Selective neuronal vulnerability in neurodegenerative diseases: from stressor thresholds to degeneration. Neuron. 2011 Jul 14;71(1):35–48. doi: 10.1016/j.neuron.2011.06.031. PMID: 21745636

Schafer, W. R. Egg-laying (December 14, 2005), WormBook, ed. The C. elegans Research Community, WormBook, doi10.1895/wormbook.1.38.1, http://www.wormbook.org.

Stefanakis N, Carrera I, Hobert O. 2015. Regulatory Logic of Pan-Neuronal Gene Expression in C. elegans. Neuron. 2015 Aug 19;87(4):733–50. doi: 10.1016/j.neuron.2015.07.031. PMID: 26291158

Taylor RC, Dillin A. 2013. XBP-1 is a cell-nonautonomous regulator of stress resistance and longevity. Cell. 2013 Jun 20;153(7):1435–47. doi: 10.1016/j.cell.2013.05.042. PMID: 23791175

Timmons L, Fire A. 1998. Specific interference by ingested dsRNA. Nature. 1998 Oct 29;395(6705):854. PMID: 9804418

Trent C, Tsuing N, Horvitz HR. 1983. Egg-laying defective mutants of the nematode Caenorhabditis elegans. Genetics. 1983 Aug;104(4):619–47. PMID: 11813735

van Oosten-Hawle P, Porter RS, Morimoto RI. 2013. Regulation of organismal proteostasis by transcellular chaperone signaling. Cell. 2013 Jun 6;153(6):1366–78. doi: 10.1016/j.cell.2013.05.015. PMID: 23746847

Ward JD. 2015. Rapid and precise engineering of the Caenorhabditis elegans genome with lethal mutation co-conversion and inactivation of NHEJ repair. Genetics. 2015 Feb;199(2):363–77. doi: 10.1534/genetics.114.172361. Epub 2014 Dec 9. PMID: 25491644

Yanik MF, Cinar H, Cinar HN, Gibby A, Chisholm AD, Jin Y, Ben-Yakar A. 2006. Nerve renegeration in Caenorhabditis elegans after femotosecond laser axotomy. IEEE Journal of Selected Topics in Quantum Electronics, Vol 12, No. 6., Nov/Dec 2006.

Yoshina S, Suehiro Y, Kage-Nakadai E, Mitani S. 2015. Locus-specific integration of extrachromosomal transgenes in C. elegans with the CRISPR/Cas9 system. Biochem Biophys Rep. 2015 Dec 1;5:70–76. doi: 10.1016/j.bbrep.2015.11.017. eCollection 2016 Mar. PMID: 28955808

Yuan J, Shaham S, Ledoux S, Ellis HM, Horvitz HR. 1993. The C. elegans cell death gene ced-3 encodes a protein similar to mammalian interleukin-1 beta-converting enzyme. Cell. 1993 Nov 19;75(4):641–52. PMID: 8242740

